# Global transcriptome analysis reveals circadian control of splicing events in *Arabidopsis thaliana*

**DOI:** 10.1101/845560

**Authors:** Andrés Romanowski, Rubén G. Schlaen, Soledad Perez-Santangelo, Estefanía Mancini, Marcelo J. Yanovsky

## Abstract

- The circadian clock of *Arabidopsis thaliana* controls many physiological and molecular processes, allowing plants to anticipate daily changes in their environment. However, developing a detailed understanding of how oscillations in mRNA levels are connected to oscillations in post-transcriptional processes, such as splicing, has remained a challenge.
- Here we applied a combined approach using deep transcriptome sequencing and bioinformatics tools to identify novel circadian regulated genes and splicing events.
- Using a stringent approach, we identified 300 intron retention, 8 exon skipping, 79 alternative 3’ splice site usage, 48 alternative 5’ splice site usage, and 350 multiple (more than one event type) annotated events under circadian regulation. We also found 7 and 721 novel alternative exonic and intronic events. Depletion of the circadian regulated splicing factor AtSPF30 homolog, resulted in the disruption of a subset of clock controlled splicing events.
- Altogether, our global circadian RNA-seq coupled with an *in silico*, event centred, splicing analysis tool offers a new approach for studying the interplay between the circadian clock and the splicing machinery at a global scale. The identification of many circadian regulated splicing events broadens our current understanding of the level of control that the circadian clock has over this posttranscriptional regulatory layer.

## INTRODUCTION

Land plants have evolved in a world that spins around its axis with a period close to 24 h. Consequently, they are subject to daily changes in the environment, such as changes in light quality / intensity and temperature. The cycling nature of these changes has allowed for the evolution of endogenous circadian timekeeping mechanisms that allow plants, and other organisms, to predict the timing of these changes, which contributes to the optimization of their growth and development (Green *et al.*, 2002; Dunlap *et al.*, 2004; Dodd *et al.*, 2005).

In *Arabidopsis thaliana*, the molecular circadian clock consists of an interlocked series of transcriptional-translational feedback loops that cycle with a period close to 24 h (Harmer *et al.*, 2001; McClung, 2014; Millar, 2016; Hernando *et al.*, 2017). The basic layout of this clock consists of the morning MYB transcription factor elements CIRCADIAN CLOCK ASSOCIATED 1 (CCA1) and LATE ELONGATED HYPOCOTYL (LHY). These elements peak in the morning and repress the transcription of TIMING OF CAB EXPRESSION 1 (TOC1), also known as PSEUDO RESPONSE REGULATOR 1 (PRR1). TOC1, in turn, peaks in the evening and represses CCA1/LHY. Another loop involves the downregulation of CCA1/LHY throughout the rest of the day by the consecutive repressive action of PRR9, PRR7 and later PRR5. During the evening, the elements EARLY FLOWERING 3 (ELF3), ELF4 and LUX ARRHYTHMO (LUX) associate into an “evening complex” (EC), repressing PRR9 and LUX. Additionally, several activators have been described. In particular, the CCA1/LHY homologs reveille 8 (RVE8), RVE6 and RVE4 are morning factors that drive the expression of PRR5 and TOC1. Together with the RVEs, the NIGHT LIGHT-INDUCIBLE AND CLOCK-REGULATED (LNK) 1-4 family of genes also contribute to activate the expression of PRR5 and ELF4 (McClung, 2014).

This circadian clock integrates environmental inputs to fine tune its oscillations (entrainment) and then controls the timing of several molecular, physiological and developmental outputs, like stomatal opening, leaf movement, hormone synthesis and growth among others (McClung, 2006; Millar, 2016). One of the many ways that this can be accomplished is through the control of gene expression, a process that involves many different steps (Romanowski & Yanovsky, 2015). One such step is the processing of precursor mRNAs (pre-mRNAs). Plant pre-mRNAs are composed of continuous nucleotide segments known as exons and introns. In order to produce a mature functional mRNA, exonic segments are kept and introns are removed through a process called splicing.

The process of splicing occurs at the spliceosome, a dynamic protein-RNA complex composed of small nuclear ribonucleoprotein particles (snRNPs) and auxiliary proteins that assemble at exon-intron boundaries known as donor 5’-splice site (ss) and acceptor 3’-ss. The selection and recognition of these splice sites occur through the interaction of spliceosome components with *cis* acting sequences and the *trans* acting proteins. Alternative splicing (AS) occurs through the alternative selection of 5’ss, 3’ss, exon inclusion or skipping and intron retention (Kornblihtt *et al.*, 2013). Through this process, different functional mRNAs can arise from the transcription of the same gene, ultimately expanding the repertoire of protein products that can be encoded by a single gene (Kornblihtt *et al.*, 2013; Saldi *et al.*, 2016).

The first evidence of crosstalk between AS and the circadian clock was described in our lab in 2010. A mutation in the clock regulated protein arginine methyltransferase 5 (PRMT5) was found to alter the AS of the clock component PRR9, causing a significant alteration in period length (Sanchez *et al.*, 2010). Since then, additional work has shown that AS is a mechanism that can couple changes in environmental temperature to circadian timing by altering the splicing of clock components (James *et al.*, 2012a; James *et al.*, 2012b; Perez-Santangelo *et al.*, 2013; Filichkin *et al.*, 2015). Also, mutations in the spliceosome components SKIP and STIPL1 can affect circadian timing (Jones *et al.*, 2012; Wang *et al.*, 2012). However, no comprehensive survey has been done to date to understand how widespread clock regulated AS is. In this work, we used mRNA-seq and a custom *in house* developed bioinformatics pipeline to perform a genome wide analysis on circadian regulation of AS in *A. thaliana* plants.

## MATERIALS AND METHODS

### Plant materials

All *Arabidopsis thaliana* plants used in this study were of the Columbia (Col-0) ecotype. *AT2G02570* mutant alleles *atspf30-1* (SALK_052016C) and *atspf30-2* (SALK_081292C) were obtained from the Arabidopsis Biological Resource Center (ABRC).

### Growth conditions and tissue collection for RNA-seq studies

Col-0 and *atspf30-1* seeds were sown on Murashige and Skoog medium (GIBCO BRL) and cold stratified (4ºC, darkness) for 3 days. For non-circadian experiments, the MS plates were then transferred to 22 °C and continuous light (LL; 50 μmol·m^−2^·s^−1^ of white light). For circadian μ experiments, the MS plates were transferred to 22 °C under LD12:12 (12 h light/12 h dark cycles; 50 μmol·m^−2^·s^−1^ of white light) conditions for 11 days and then transferred to continuous light (LL; 50 μmol·m^−2^·s^−1^ of white light).

Samples containing 15-30 seedlings were flash frozen in liquid nitrogen. For circadian studies, the samples were collected at the timepoints indicated in Fig. 1A, with 2 biological replicates per timepoint. For non-circadian studies, 3 biological replicates of Col-0 and *atspf30-1* were sampled. After all samples were collected, total RNA was extracted using the Trizol method according to the manufacturer’s protocol (ThermoFisher) and quantified using a NanoDrop spectrophotometer (ThermoFisher). Total RNA integrity was further checked by agarose gel electrophoresis.

**Fig. 1.**
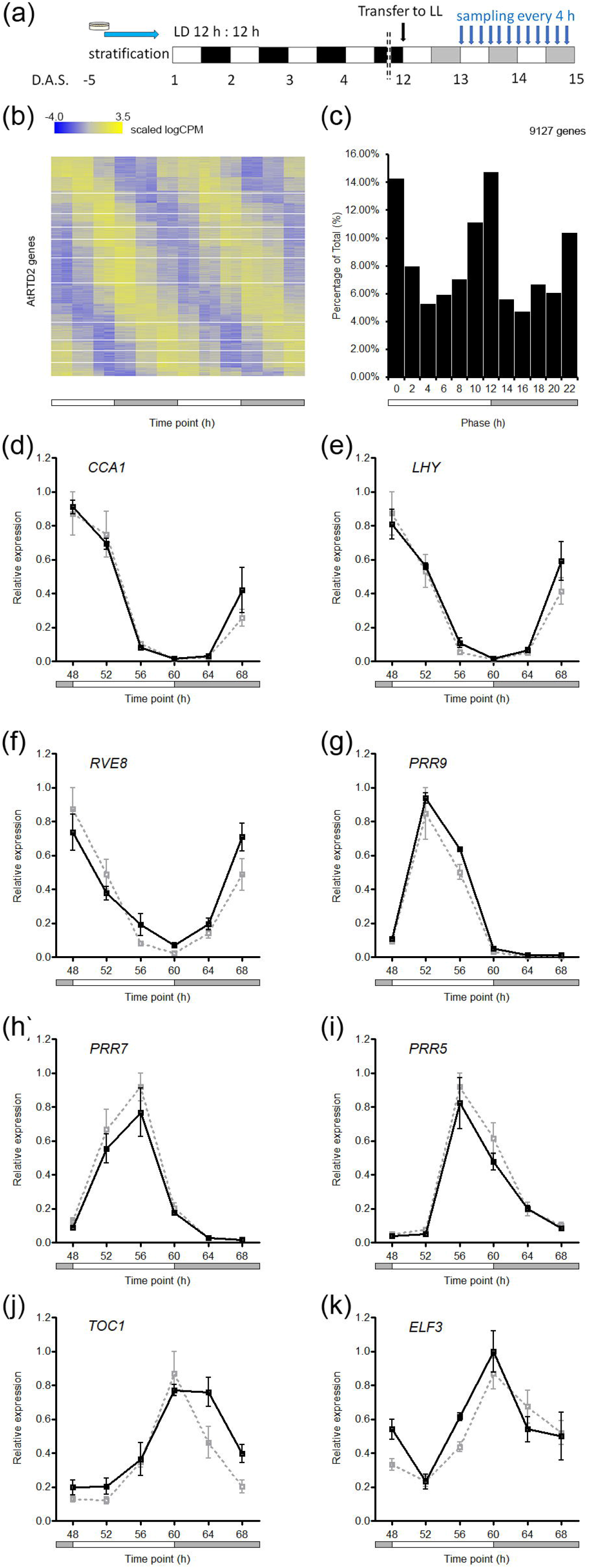
Global circadian gene expression. (a) Schematic of experimental conditions. (b) Phase sorted heatmap of global scaled logCPM gene expression from the circadian timecourse experiment. (c) Phase histogram of global gene expression. (d-k) RT-qPCR validation (black, solid line) and RNA-seq traces (grey, dotted line) of the core clock genes (d) CCA1, (e) LHY, (f) RVE8, (g) PRR9, (h) PRR7, (i) PRR5, (j) TOC1, and (k) ELF3. Expression values are normalised to maximum. Error bars represent SEM. CPM: counts per million. CCA1: Clock Control Associated 1 (AT2G46830). LHY: Late Elongated Hypocotyl (AT1G01060). RVE8: Reveille 8 (AT3G09600). PRR9: Pseudo Response Regulator 9 (AT2G46790). PRR7: Pseudo Response Regulator 7 (AT5G02810). PRR5: Pseudo Response Regulator 5 (AT5G24470). TOC1: Timing of CAB Expression 1 (AT5G61380). ELF3: Early Flowering 3 (AT2G25930).

### Library preparation and RNA sequencing

Libraries were prepared following the TruSeq RNA v2 Sample Preparation Guide (Illumina). Briefly, 3 μg of total RNA was polyA-purified and fragmented, and first-strand cDNA synthesized by reverse transcriptase (SuperScript II; Invitrogen) and random hexamers. This was followed by RNA degradation and second-strand cDNA synthesis. End repair process and addition of a single A nucleotide to the 3’ ends allowed ligation of multiple indexing adapters. Then, an enrichment step of 12 cycles of PCR was performed. Library validation included size and purity assessment with the Agilent 2100 Bioanalyzer and the Agilent DNA 1000 kit (Agilent Technologies). Samples were diluted to 10nM and pooled to create the 12X multiplexed DNA libraries, and 12–15 pmol of material was loaded onto each lane. Template hybridization, extension of the template, amplification of the template, cluster generation, sequencing primer addition, and single end chemistry were performed using C-BOT (formerly known as Cluster Station). Base calls were made using CASAVA.

The multiplexed samples were loaded onto two lanes on an Illumina HiSeq 2500 sequencer, providing 100-bp single-end reads.

The RNA-seq fastq files supporting the conclusions of this article are available in the ArrayExpress (Kolesnikov *et al.*, 2015) database at EMBL-EBI (www.ebi.ac.uk/arrayexpress), under accession numbers E-MTAB-7933 (circadian dataset) and E-MTAB-8667 (atspf30-1 dataset).

### Processing of RNA reads and alignment

Reads were quality-filtered using the standard Illumina process and demultiplexed with two allowed barcode mismatches. Sequence files were generated in FASTQ format. The Bowtie/TopHat suite (Trapnell *et al.*, 2009) was used to map reads to the *A. thaliana* TAIR10 reference genome. Along with the prebuilt *A. thaliana* index, the reference genome was downloaded from ENSEMBL (December 2014). Default values for TopHat parameters were used with the exception of maximum intron length parameter, which was set to a value of 5,000 nt following estimated values, as previously reported (Hong *et al.*, 2006). Table S1 provides a summary table of main read count statistics.

### RNA sequencing analysis

Read counts were calculated for genes, exon bins, AS bins and intron bins with the ASpli package (Mancini *et al.*, 2019) using custom scripts written in R (64-bit, version 3.6.0). Multidimensional scaling (MDS) plots of distances between gene expression profiles were generated with the limma package (Ritchie *et al.*, 2015), using custom script written in R. Coverage plots were generated using the IGB Browser (Freese *et al.*, 2016).

Detection of cycling was performed using JTK_Cycle (Hughes *et al.*, 2010) implemented in R, using 24±4 h periods as limits. Obtained p-values were FDR corrected. Recalculated circadian phases were obtained by multiplying the JTK obtained phase (LAG) by 24 and dividing that number by the JTK obtained period (PER). Relative amplitude estimates were calculated by dividing the subtraction of the absolute peak expression and through levels, by the sum of through level plus one (to avoid divisions by zero). Heatmaps were generated using scripts for R. PSI/PIR indexes were calculated as previously described (Pervouchine *et al.*, 2013; Gueroussov *et al.*, 2015) using ASpli implemented in R. GO analyses were performed with the Virtual Plant web service (Katari *et al.*, 2010). All code and analyses are available on demand.

### Semiquantitative and reverse transcriptase PCR (sqRT-PCR)

Plants were grown and sampled as described above (see ‘Growth conditions and tissue collection for RNA-seq studies’). Total RNA was obtained from these samples using TRIzol reagent (Invitrogen). One microgram of RNA was treated with RQ1 RNase-Free DNase (Promega) and subjected to retro transcription with Superscript III (Invitrogen) and oligo dT according to the manufacturer’s instructions. Amplification of cDNA was carried out for 30 cycles to measure splicing at the linear phase. RT-PCR products were electrophoresed in 8% SDS-PAGE gels and detected with EtBr. Primers used for amplification of each gene are described in Table S2.

### Quantitative reverse transcriptase PCR (qRT-PCR)

cDNAs obtained for ‘Semiquantitative and reverse transcriptase PCR (sqRT-PCR)’ were also used for quantitative RT-PCR amplification. Synthesized cDNAs were amplified with FastStart Universal SYBR GreenMaster (Roche) using the Mx3000P Real Time PCR System (Agilent Technologies) cycler. The PP2A (AT1G13320) transcript was used as a reference gene.

Quantitative RT-PCR quantification was conducted using the standard curve method as described in the Methods and Applications Guide from Agilent Technologies. Primers and targets for gene expression and splicing events validations are described in Table S2.

### Circadian Leaf Movement Analysis

For leaf movement analysis, Col-0 and *atspf30-1* plants were entrained under a 16 h light: 8 h dark cycle, transferred to continuous white fluorescent light, and the position of the first pair of leaves was recorded every 2 h for 5–6 d using Image J software (http://imagej.nih.gov). Period estimates were calculated using Brass 3.0 software (Biological Rhythms Analysis Software System; http://www.amillar.org). Briefly, raw data was imported; baseline detrended and analysed using the FFT-NLLS algorithm, as described previously (Plautz *et al.*, 1997). Statistical analysis was performed using a two tailed Student’s t test, after testing for normal distribution of the data.

### Flowering Time Analysis

For flowering time analysis, Col-0, *atspf30-1* and *atspf30-2* plants were grown under long days (16 h light: 8 h dark), short days (8 h light: 16 h dark) or continuous light at a constant temperature of 22 °C. Flowering time was estimated by counting the number of rosette leaves at the time of bolting. Statistical analysis was performed using Dunnet’s multiple comparisons test using Col-0 as a control, after testing for normal distribution of the data.

### Phylogenetic Tree Reconstruction

Splicing Factor 30 (SPF30) and Survival Motor Neuron (SMN) protein sequences from *Homo sapiens, Rattus norvergicus, Mus musculus, Danio rerio, C. elegans* and *D. melanogaster.* Homolog sequences from *A. thaliana, Capsella rubella, Solanum lycopersicum, Oryza sativa, Sorghum bicolor*, were identified with BlastP and downloaded from GenBank. *Physcomitrella patens* sequences were obtained from PEATMoss PpGML DB v1.6 (Fernandez-Pozo *et al.*, 2019). Using these sequences as input, a phylogenetic tree was reconstructed using SeaView 4.7 (Gouy *et al.*, 2010). Sequences were aligned with Clustal Omega 1.2.0 (Sievers *et al.*, 2011) with default options. PhyML 3.1 (Guindon *et al.*, 2010) was used for the phylogenetic tree reconstruction, using default parameters except model was set to LG amino-acid replacement matrix (Le & Gascuel, 2008), and bootstrapping was set to 1000 replicates.

## RESULTS

### Global RNA-seq of the circadian transcriptome reveals known and novel circadian regulated genes

To analyse the circadian regulation of gene expression, we performed a global transcriptome deep sequencing of 15-day old *A. thaliana* plants grown under a light/dark (LD) 12 h: 12 h photoperiod, transferred to constant light conditions (LL), and harvested after 24 h in LL every 4 h for a total of 2 days, with two biological replicates (Fig. 1a). This approach, in contrast to previous studies that used microarray-based approaches, allowed us to analyse genes which are not represented in microarrays. The data generated here represent more than 361 million short nucleotide reads (100bp length) and 36.1 Gb of successfully aligned sequence (Table S1). This constitutes the deepest sequencing of a plant circadian transcriptome to date.

Gene expression was computed with the ASpli R package (Mancini *et al.*, 2019) using the AtRTD2 annotation (Zhang *et al.*, 2017) by obtaining a table of gene counts, which represent the total number of reads aligning to each gene (34,212 annotated AtRTD2 genes). On average, the reproducibility between replicate samples (i.e., samples from the same time point) was excellent, with Pearson correlation average R^2^ values approaching 0.99 (see Supplemental Fig. 1 online). Using a Multidimensional scaling (MDS) plot we also verified that samples corresponding to the same timepoint on the two consecutive free running days (LL day 2 and LL day 3) were similar (Fig. S1). We then obtained the number of reads per gene length (read density) for each gene, and only those that had a value higher than 0.05 in any of the time points throughout the time course were chosen for further analysis (18,503 genes).

In order to improve the power of subsequent rhythmic statistical tests, we applied a pre-screening step to exclude any obviously non-rhythmic transcripts, as previously suggested (Keegan *et al.*, 2007). Only the genes that showed a variation throughout the time-course (13,256 genes; negative binomial generalized linear model (GLM) model fit, p<0.05 and q<0.10) were then subjected to circadian rhythmicity analysis. To this end, several statistical approaches have been described (Wichert *et al.*, 2004; Wijnen *et al.*, 2005; Michael *et al.*, 2008; Deckard *et al.*, 2013). In this work, we applied the Jonckheere-Terpstra-Kendall (JTK) algorithm, which has been reported to identify rhythmic genes with enhanced confidence, improved computational efficiency and more resistance to outliers than other algorithms (Hughes *et al.*, 2010). Using this method, a total of 9,127 genes were found to be rhythmic and exhibited a circadian expression pattern (JTK_cycle, p<0.01 and q<0.01; Fig. 1b, Table S3) under our assay conditions. To verify the validity of our approach, we examined the expression pattern of known core clock genes in our data set and validated them by qPCR (Fig. 1d-k). As expected, the qPCR reproduced the patterns observed on our RNA-seq data (Fig. 1d-k). As expected for circadian datasets, peak amplitude on day 3 is slightly lower than peak amplitude on day 2, indicating a progressive decrease of amplitude in constant conditions (Table S3).

We then performed a Gene Ontology (GO) analysis of this dataset (Fig. S2, Table S4) which showed that clock controlled genes (CCGs) were involved in diverse biological processes. These included, but were not limited to, immune response, hormone signalling, light signalling, metabolism and photosynthesis, which is in accordance to previous reports (Covington *et al.*, 2008; Hsu & Harmer, 2012). Amplitude histogram analysis reveals that most transcripts oscillate with a relative amplitude below 3, while a minority of circadian transcripts oscillate at high amplitude. This latter set includes the core circadian clock genes (Fig. S3), and is consistent to what is known in mammals and fruit flies (Hughes *et al.*, 2009; Hughes *et al.*, 2012).

A phase distribution analysis of the circadian regulated genes showed that that 25.9% (2364 genes) exhibit a peak phase expression at LL10-12, while 24.66% (2251 genes) exhibit a peak phase expression at LL22-LL0. These correspond to the day-to-night and night-to-day transitions of the LD 12 h: 12 h entraining photoperiod, respectively (Fig. 1b). Genes that peaked at LL10-12 of the free running cycle were involved in 96 biological processes (Fisher’s Exact Test, p < 0.05), including response to stress, immune response, lipid metabolism, signalling and protein processes (Fig. S4a). Genes that peaked at LL22-0 were involved in 146 biological processes (Fisher’s Exact Test, p < 0.05), including RNA modification/processing, glucosinolate biosynthesis, carbohydrate metabolism, mitochondrial/intracellular transport and ribosome biogenesis (Fig. S4b)

Upon comparison of our dataset with previous published global *A. thaliana* circadian gene expression datasets (Covington *et al.*, 2008; Hsu & Harmer, 2012) (Table S5), we found an overlap of 41.67% (3,803 genes) and 49.76% (4,542 genes) to the Covington + Edwards combined gene lists (Covington *et al.*, 2008) (DS1) and Hsu (Hsu & Harmer, 2012) (DS2) datasets, respectively. A total of 2,763 genes (30.27%) were found to cycle in all three datasets (Fig. 2a). By combining the three datasets, we find that over 37.29% of *A. thaliana* genes (12,579 / 34,212 AtRTD2 genes) have been described to be circadian regulated to this date, in global transcriptomic studies (Table S5).

**Fig. 2.**
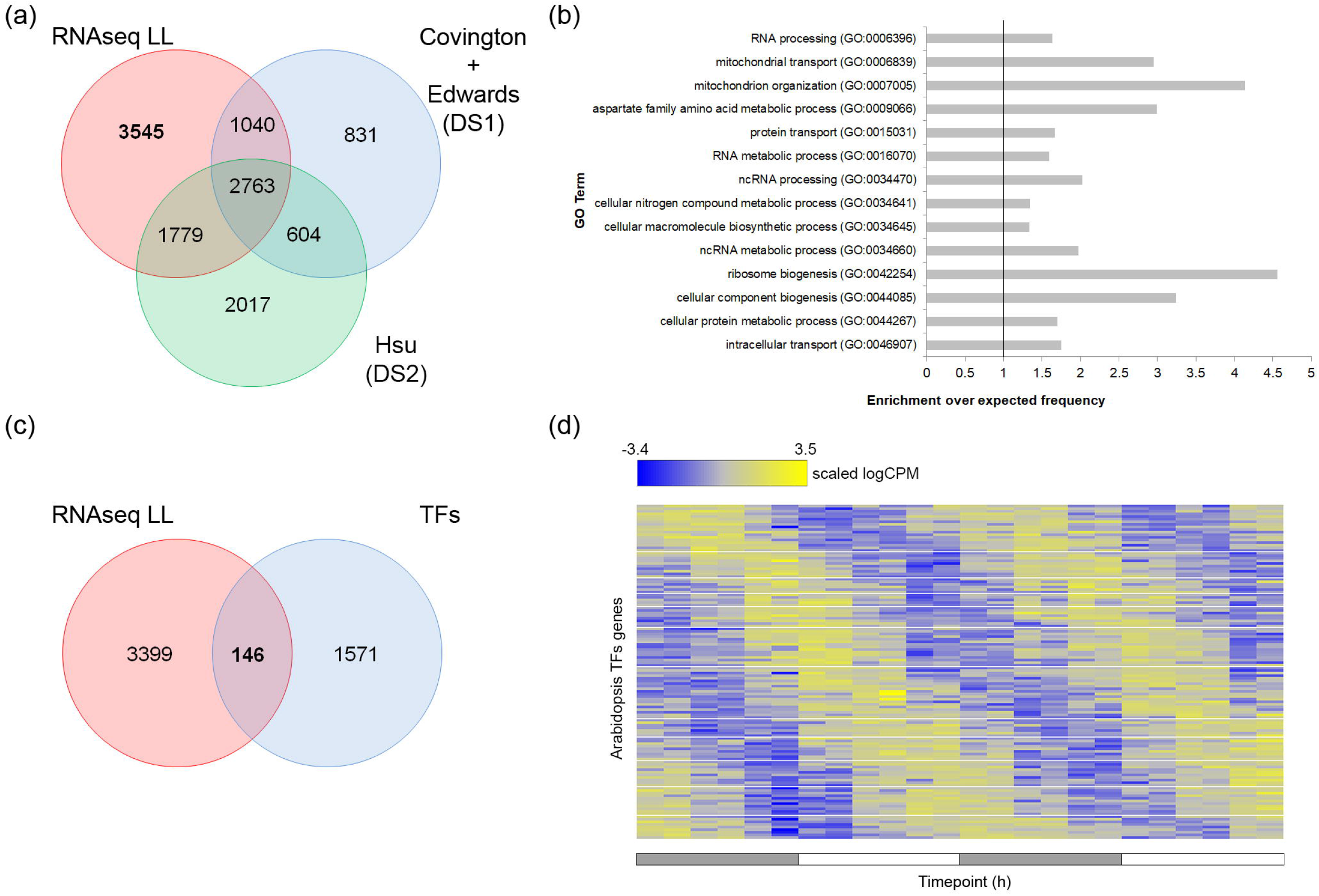
Novel circadian genes and specific gene families. (a) Venn diagram showing RNA-seq circadian gene expression (RNA-seq LL) compared to the Covington + Edwards combined dataset (DS1) (Covington *et al.*, 2008) and the Hsu & Harmer dataset (DS2) (Hsu & Harmer, 2012). (b) Selected subset of significant Biological Process (BP) GO Terms of novel circadian genes. Black line indicates enrichment above expected frequency. (c-d) Novel cycling transcription factor genes. (c) Venn diagram showing the comparison between the novel genes of the circadian dataset and all known transcription factor genes. (d) Phase sorted heatmap of scaled logCPM gene expression of the 146 novel cycling transcription factor genes. DS1: Dataset 1. DS2: Dataset 2. CPM: Counts per million.

### Novel circadian transcripts

Our dataset also revealed novel circadian regulated genes that had not been identified by the two previous global studies (Fig. 2a). This was to be expected, to a certain degree, because of the difference in genome coverage of the different technologies used (RNA-seq, ATH1 microarray and AGRONOMICS1 tiling array). However, hypothetically there should be no difference in the possible number of positive genes between our RNA-seq approach and the AGRONOMICS1 platform (Rehrauer *et al.*, 2010) (Fig. S5). Nevertheless, we found 3545 circadian genes that are absent from previous global studies. We will refer to these genes as novel circadian genes. GO analysis of this subset revealed an enrichment in terms related to translation, ribosome biogenesis and RNA metabolism (Fig. 2b, Table S6). It should be noted that some of the genes in this subset have been previously identified as circadian regulated in non-global studies. Such is the case, for example, of the widely used circadian reporter *CAB2* (Millar & Kay, 1991).

Interestingly, 146 transcription factors (TFs) spanning many families, including bZIP, bHLH, MADS box related, WRKY, MYB and NAC, are part of the novel CCGs subset (Fig. 2c, Table S3). The RNA-seq data of the expression pattern of these TFs genes can be seen in Fig. 2d. Further study of these genes might lead to new insight into how the circadian clock regulates gene expression.

### The circadian clock regulates alternative splicing events

In this work, we took advantage of the accuracy of RNA-seq technology and then applied a custom pipeline designed to specifically detect and quantify splicing events. To do so, we aligned the RNA-seq reads to the TAIR10 reference genome to generate the BAM files required as input for the *in house* developed ASpli Bioconductor R package. ASpli was designed by our lab for the specific purpose of detecting and quantifying splicing events (Mancini *et al.*, 2016; Mancini *et al.*, 2019). This package has successfully been employed to study alternative splicing in several studies spanning different organisms (De Maio *et al.*, 2016; Mancini *et al.*, 2016; Beckwith *et al.*, 2017; Xin *et al.*, 2017). In summary, when used to detect changes in splicing events, this algorithm uses a known genome annotation to categorise the genome in discrete exon and intron “bins”, and then quantifies the counts that each bin has over time and normalizes it to gene expression. After obtaining the normalized bin counts, we proceeded to analyse the rhythmicity of splicing events as explained for gene expression (Mancini *et al.*, 2016; Mancini *et al.*, 2019). Using this method, coupled to the AtRTD2 annotation (Zhang *et al.*, 2017), 3976 clock controlled splicing events (CCEs) were discovered. In particular, 300 intron retention (IR), 40 exon skipping, 78 alternative 5’ splice site (Alt5’ss), 113 alternative 3’ splice site (Alt3’ss), 470 multiple (involved simultaneously in more than one AS event type), 22 undefined, and 1720 novel exonic and 1233 novel intronic events were found to be rhythmic, and exhibited a circadian expression pattern (JTK_cycle, p<0.01 and q<0.01) with relative amplitude > 0.15 under our assay conditions. With the purpose of increasing the stringency of our approach, we applied an extra filtering step and kept only those splicing events with evidence of splice-junction reads. To do so, we calculated the Percentage of Spliced In (PSI) and the Percentage of Intron Retention (PIR) indexes as previously described (Gueroussov *et al.*, 2015) for exon and intron bins, respectively. We kept the subset of circadian splicing events that showed more than a 5% variation in PSI/PIR (dPSI/PIR) across all the analysed timepoints. This resulted in a total of 1513 events corresponding to 1169 genes, split into the following categories: 79 Alt3’ss, 48 Alt5’ss, 300 IR, 8 ES, 350 multiple (Fig. 3a), and 7 novel exonic and 721 novel intronic (Fig. 3b) (Table S7). It is interesting to note that this study also revealed that 617 CCGs are also clock regulated at the level of alternative use of exons and/or introns throughout the day (Fig. S6a, Table S7). On the other hand, there also are 552 genes whose expression is not circadian regulated, but their exon/intron use is (Fig. S6a, Table S7), and 8510 CCGs that had no detectable CCEs. In total, 652 genes had novel exonic/intronic events, 69 Alt3’ss, 48 Alt5’ss, 286 IR, 8 ES and 292 multiple (Fig. S6b), including several splicing factors and spliceosome components.

**Fig. 3.**
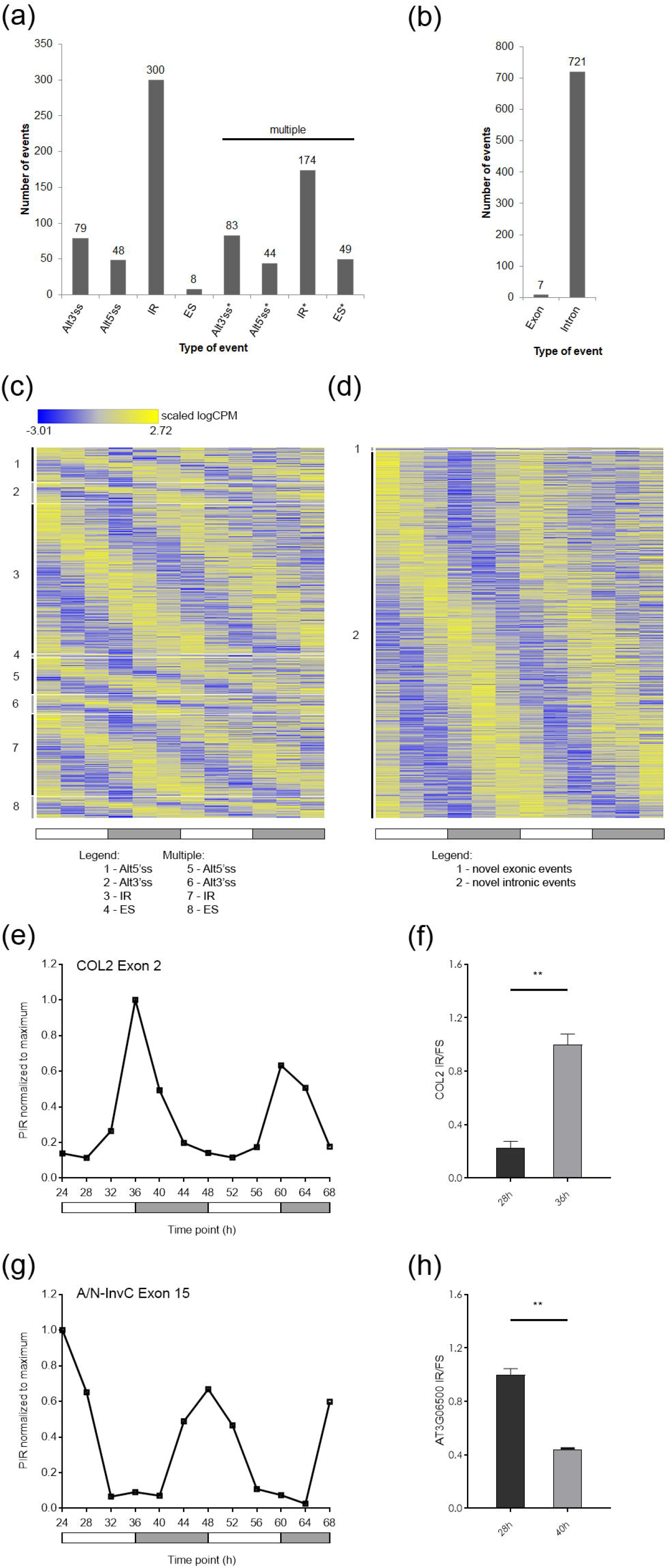
Analysis of Clock Controlled Elements (CCEs). (a) Distribution of clock controlled annotated events by type (alt 3’ss, alt 5’ss, ES, IR and multiple). (b) Distribution of novel (non-annotated) clock controlled exonic and intronic events. (c-d) Phase sorted heatmap of scaled logCPM clock controlled splicing event bins expression by type: (c) annotated events (alt 3’ss, alt 5’ss, ES, IR and multiple), (d) non-annotated events (exonic and intronic). (e-f) Validation of CCEs. (e) PIR normalised to maximum of COL2 E2. (f) qPCR validation of the COL2 E2 IR event (** p<0.01; Student’s T-test). (g) PIR normalised to maximum of the A/N-INvC E15. (h) qPCR validation of the A/N-INvC E15 IR event (** p<0.01; Student’s T-test). Error bars represent SEM. alt 3’ss: alternative 3’ splice site. alt 5’ss: alternative 5’ splice site. ES: exon skipping. IR: intron retention. CPM: counts per million. CCEs: clock controlled splicing events. PIR: Percent Intron Retention. COL2: CONSTANS like 2 (AT3G02380). A/N-INvC: Alkaline/Neutral Invertase C (AT3G06500).

We then performed a Gene Ontology (GO) analysis of this dataset (Table S8) which showed that clock controlled splicing events (CCEs) included genes involved in primary metabolism (GO:0044283), nitrogen metabolism (GO:0006807 and GO:0034641), nucleic acid metabolism (GO:0006139 and GO:0090304), RNA/mRNA processing (GO:0016070, GO:0006396, GO:0006397, GO:0016071), lipid metabolism (GO:0006644, GO:0009245, GO:0046493 and GO:0008654), flowering (GO:0009909 and GO:0009910), and RNA splicing (GO:0000375, GO:0000377, GO:0008380).

In order to verify the validity of our approach, we performed an independent experiment and examined the expression pattern of a couple of splicing events from our data set and validated them by semi-quantitative (sq)RT-PCR, followed by SDS-PAGE. Results were then further corroborated by qPCR, obtaining in both cases results that matched the patterns produced by our RNA-seq analysis pipeline data (Fig. 3e-h, Fig. S7a-b). We validated the intron retention splicing event of the COL2 gene (IR - AT3G02380:E002 – width 143bp, Table S7), homolog of the photoperiod and flowering regulator CONSTANS. This event had been previously described (Hazen *et al.*, 2009) and was therefore a perfect positive control. According to the bioinformatics analysis of our pipeline, the IR precedes the subjective day and has a peak at the day/night transition, as can be seen in the RNA-seq coverage plots (Fig. S7a) and PIR pattern (Fig. 3e). This event also fulfils our stringent criteria of having reads that span the intron-exon. The analysis by sqRT-PCR followed by SDS-PAGE (Fig. S7b) and qPCR (Fig. 3f) demonstrate the validity of the bioinformatics analysis used in this work. Additionally, our results exhibit a peak IR at CT36, which is in complete accordance to what was previously described (Hazen *et al.*, 2009), and further validates our analysis pipeline. Additionally, we also validated an IR event on the A/N-InvC gene (IR - AT3G06500:E015 – width 81bp, Fig. 3g-h). This gene is expressed in roots, aerial parts (shoots and leaves) and flowers, and encodes an alkaline/neutral invertase which localizes in mitochondria. Mutations of this gene have been shown to have reduced shoot growth (Martin *et al.*, 2013). Retention of this intron causes inclusion of an in-frame Opal stop codon (at position 542), leading to a truncated protein with 123 less amino acid residues.

### Several splicing modulators are under circadian clock control

We reasoned that this circadian control of alternative splicing could be due, at least in part, to an underlying circadian regulation of the splicing machinery. To test this hypothesis, we compiled a list of all known splicing modulators, as previously described (Perez-Santangelo *et al.*, 2013). This list contains includes SR, hnRNPs, core spliceosomal proteins, regulators of spliceosome assembly, components of specific snRNPs and some kinases that regulate SR protein function. We then intersected it with the Covington + Edwards dataset (DS1), the Hsu & Harmer dataset (DS2) and our own. Out of the 426 splicing related genes, 31 were rhythmic in all three datasets (Fig. 4a). Because different research groups, each with their own environmental conditions, obtained these datasets independently, we reasoned that these 31 splicing modulators would be the most robustly rhythmic ones.

**Fig. 4.**
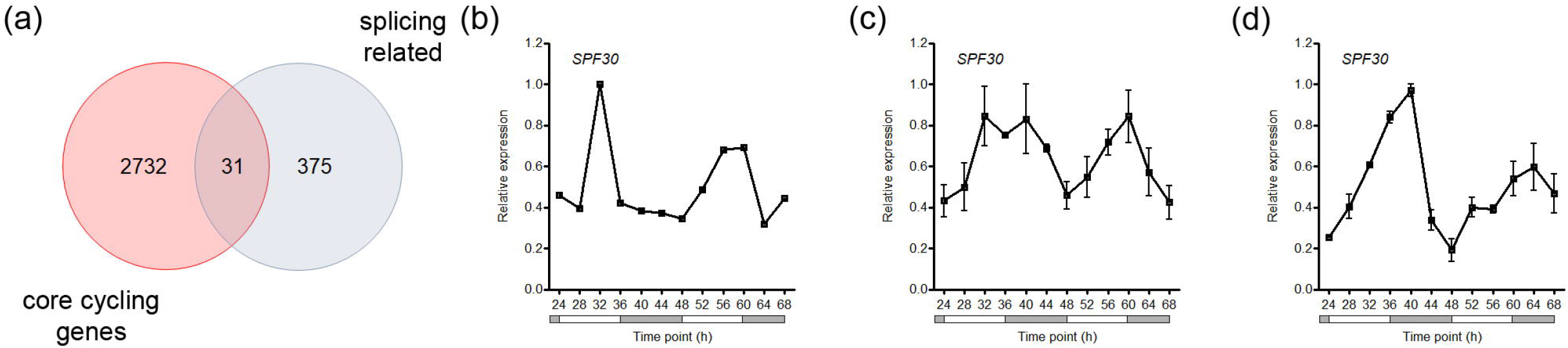
Circadian regulated splicing factors and AtSPF30. (a) Venn diagram showing the comparison between robust cycling genes (those that cycle on the RNA-seq dataset of this work and also on DS1 and DS2) and splicing related genes. (b-d) AtSPF30 expression pattern from: (b) Mockler Diurnal web server, LL23_LDHH dataset (Mockler *et al.*, 2007), (c) Normalised counts from the circadian RNA-seq (this work), and (d) qPCR data (n=3 biological replicates per timepoint). Expression values are normalised to maximum. Error bars represent SEM. AtSPF30: Arabidopsis thaliana Splicing Factor 30 (AT2G02570).

One of these genes, AT2G02570, has been previously described as the homolog of the mammalian Survival of Motor Neuron (SMN), which controls spliceosome assembly (Kroiss *et al.*, 2008). However, our reciprocal BlastP search revealed that AT2G02570 is more closely related to Splicing Factor 30 (SPF30, e.g. NP_005862.1) (Fig. S8). SPF30 is an SMN paralogue required for spliceosome assembly, which as SMN also contains a Tudor domain (Cote & Richard, 2005; Liu *et al.*, 2012; Mier & Perez-Pulido, 2012). This domain has affinity to Sm ribonucleoproteins that form a ring involved in the splicing process (Cote & Richard, 2005). Based on increasing evidence indicating that alternative splicing is regulated not only by SR and hnRNP auxiliary factors, but also by changes in the levels or activities of core spliceosomal components or proteins that modulate the kinetics of spliceosome assembly (Saltzman *et al.*, 2011; Papasaikas *et al.*, 2015), we decided to further investigate the biological relevance of AT2G02570, which will be called AtSPF30 throughout the text.

### AtSPF30 affects a subset of clock-regulated events

In order to investigate the role of AtSPF30 on the modulation of CCEs, we first validated its rhythmic transcriptional pattern by qPCR (Fig. 4b-d) and isolated two different T-DNA insertion mutants (*atspf30-1* and *atspf30-2*, Fig. 5a). Plants homozygous for either mutant allele were morphologically very similar to the wild type (Fig. 5a). Characterization of two clock regulated outputs, leaf movement and flowering, revealed no significant difference in leaf movement periodicity between Col-0 and either mutant (Fig. 5b) and a consistent slightly early flowering phenotype across the different photoperiods tested (LL, LD and SD, *p<0.05 ANOVA followed by Dunnett’s *post hoc* test against Col-0; Fig. 5c).

**Fig. 5.**
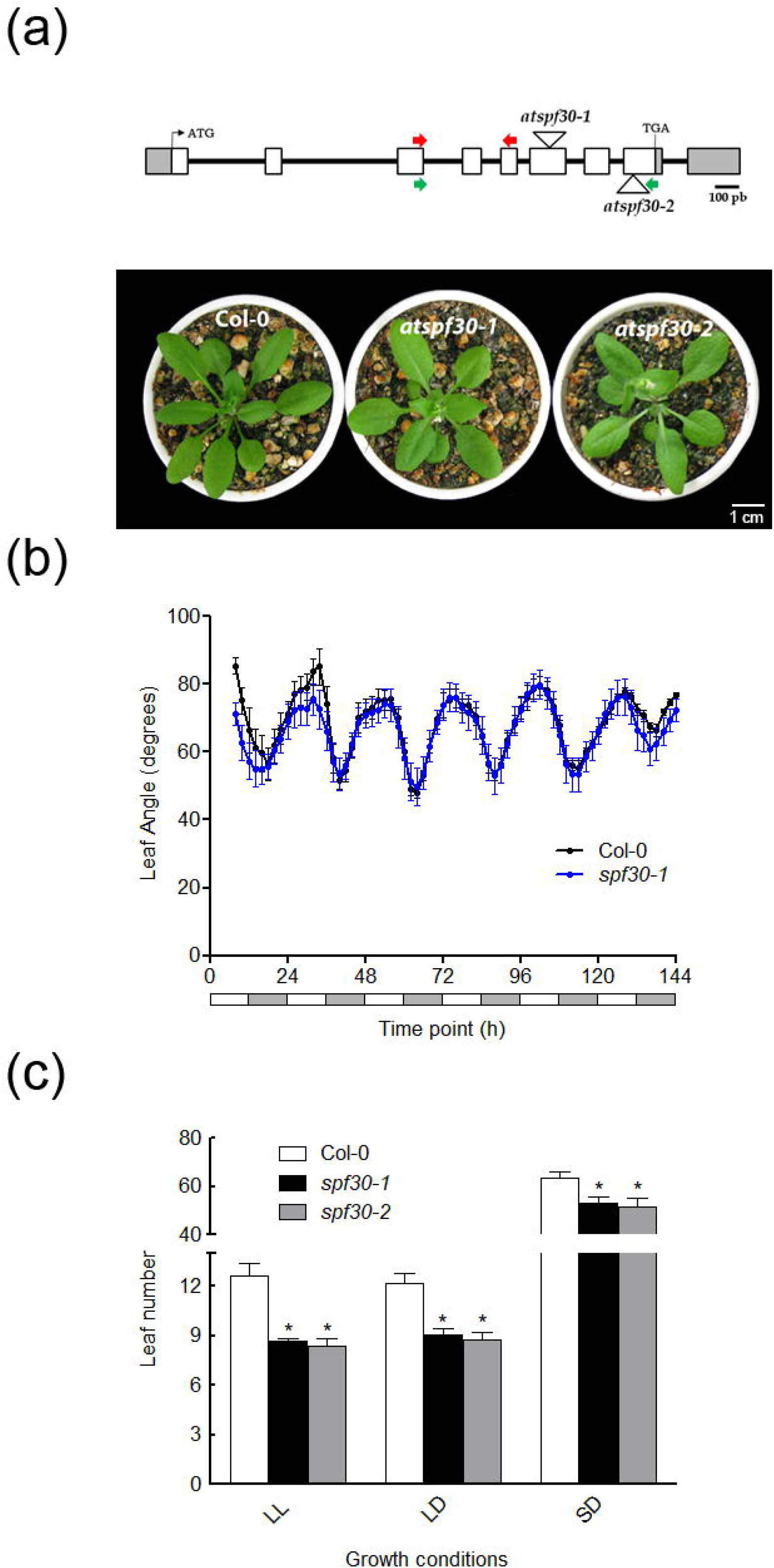
AtSPF30 alleles, circadian leaf movement and flowering phenotype. (a) Gene model of AtSPF30. White triangles indicate the T-DNA insertion sites for the mutant alleles *atspf30-1* (SALK_052016C) and *atspf30-2* (SALK_081292C). Red and green arrows indicate primer binding sites. Representative images of wild type (Col-0), atspf30-1 and atspf30-2 (from left to right) are shown. (b) Graph showing circadian leaf angle plots for Col-0 (black) and atspf30-1 (blue), after transfer to LL for 144 h (6 days). These experiments were repeated at least three times with similar results. No differences in period were found among genotypes, as estimated by Fast Fourier Transform-Non-Linear Least Squares (FFT-NLLS) algorithm with BRASS 3.0. software. (c) Flowering time measured as the number of rosette leaves at 1 cm bolting in LL, SD and LD photoperiods (n = 3 independent experiments for LL and LD, and n = 4 independent experiments for SD). Error bars represent SEM. * p<0.05 (ANOVA, followed by Dunnet’s *post hoc* test against Col-0). AtSPF30: Arabidopsis thaliana Splicing Factor 30 (AT2G02570). LL: continuous light. SD: short days. LD: long days.

RNA-seq analysis of *atspf30-1* mutant plants grown in constant light revealed 286 altered alternative splicing events when compared to Col-0 plants grown in the same conditions (|FC| > 1.5, p<0.05, q<0.05 and dPSI/PIR > 5%; Fig. 6a, Table S9). In wild-type plants, the most abundant AS events were those associated with the use of alternative 3 splice sites (Alt3’ss; 33%), followed by intron retention (IR; 32%), alternative 5 splice sites (Alt5’ss; 22%), and exon skipping (ES; 13%). However, among the AS events altered in the *atspf30-1* mutant, we found an increase in the proportion of IR events, a decrease in 3’alt events and 5’alt events relative to their frequency in wild type plants (Fig. 6a-b). Interestingly, 32 of these events were also among our PSI/PIR filtered subset of CCEs (Fig. 6c, Table S10). Upon closer inspection of the dPSI/PIR of *spf30-1* vs Col-0 of the 32 events in common between both datasets (Table S10), we hypothesised that intron retention above the level of wild type would also be expected on *atspf30-1* along an entire circadian period. With this in mind, we selected an IR event that was both, affected by a mutation in AtSPF30 and clock controlled, with a peak phase at CT16, for validation (Fig. 6d-f). qPCR quantification of a new set of circadian timecourse samples showed that lack of functional AtSPF30 affected the overall inclusion of the of AT5G17670:E006 along the entire circadian day without affecting the rhythmic pattern (Fig. 6f). Hence, in the case of this splicing event, AtSPF30 is modulating both the baseline level of intron retention and its overall amplitude.

**Fig. 6.**
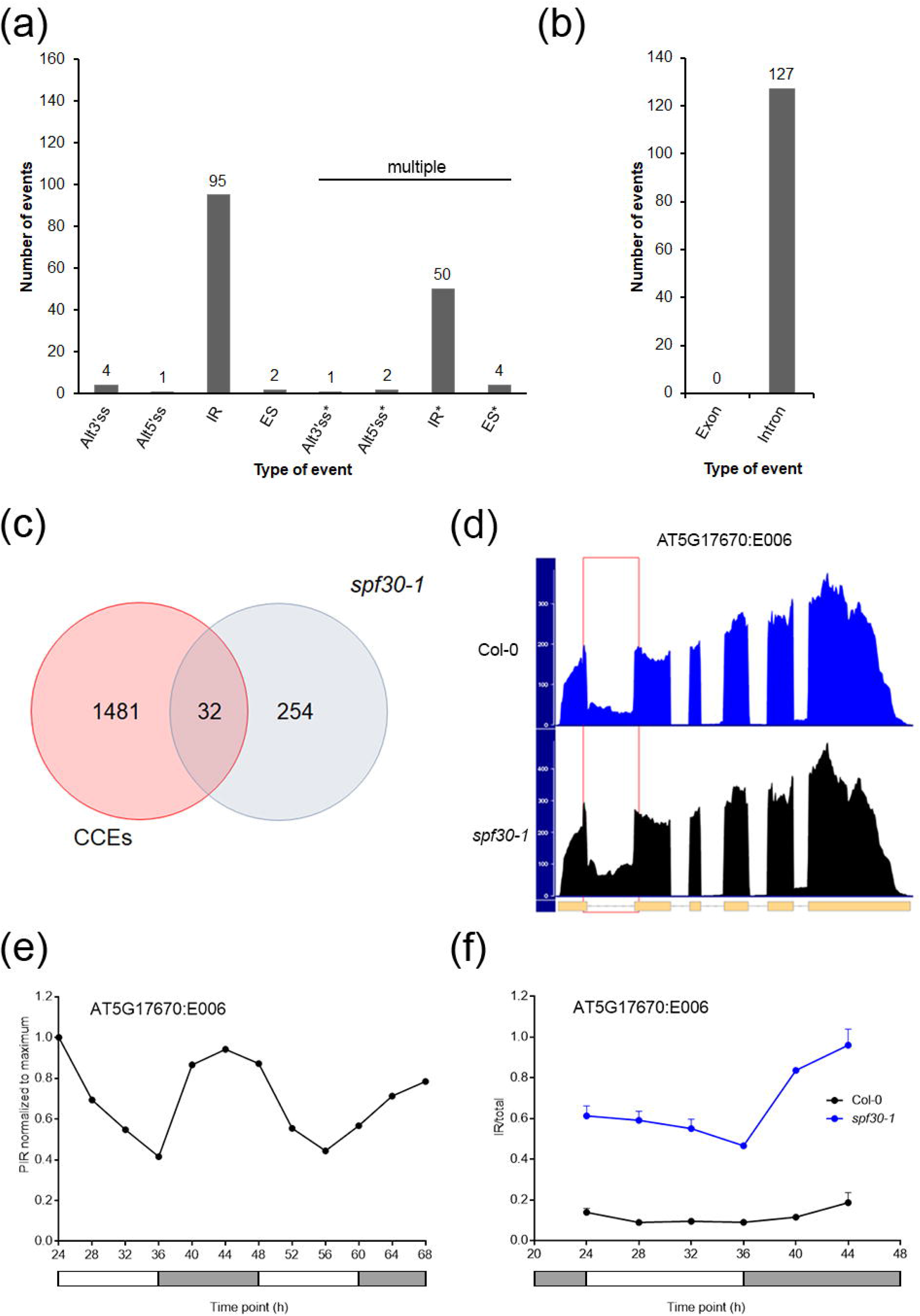
CCEs misregulated in atspf30-1. (a-b) Distribution of events affected in atspf30-1 plants. (a) Annotated events by type (alt 3’ss, alt 5’ss, ES and IR) (b) Non-annotated events (exonic and intronic). (c) Venn diagram showing the comparison between all clock-controlled elements (CCEs) and atspf30-1 affected splicing events. (d) Read density histogram of RNA-seq data comparing Col-0 (blue) and atspf30-1 (black) reads for the AT5G17670 gene locus region. The red rectangle area depicts the read density zone for the IR event. The AT5G17670 gene model is shown at the bottom of the graph. (e) PIR plot of the AT5G17670:E006 event. (f) qPCR of AT5G17670:E006 in Col-0 (black) and atspf30-1 (blue). Expression values are normalised to maximum. Error bars represent SEM. alt 3’ss: alternative 3’ splice site. alt 5’ss: alternative 5’ splice site. ES: exon skipping. IR: intron retention. PIR: percent intron retention.

## DISCUSSION

AS is highly prevalent in plants, affecting pre-mRNAs of more than 60% of intron-containing genes (Reddy *et al.*, 2013). Many genes undergo AS in response to development and changes in the environment, such as in light and temperature (James *et al.*, 2012a; James *et al.*, 2012b; Reddy *et al.*, 2013; Kwon *et al.*, 2014). For some time now, evidence has been accumulating pointing to the fact that AS can affect clock components and that mutations affecting spliceosome related components can alter the circadian period (Sanchez *et al.*, 2010; Jones *et al.*, 2012; Perez-Santangelo *et al.*, 2013; Cui *et al.*, 2014; Filichkin *et al.*, 2015). Furthermore, studies performed in fruit flies, mice and human cell lines have identified a role for the circadian clock in the control of the splicing process (Hughes *et al.*, 2012; Preussner *et al.*, 2017; Genov *et al.*, 2019). However, there have been no global transcriptomic studies that allowed for understanding the extent of clock control on AS on any plant species. There is perhaps one interesting exception: a global study limited to the analysis of cycling introns using GeneChip AtTILE1 tiling array technology (Hazen *et al.*, 2009). A total of 499 genes were reported to contain cycling introns at the time. In this study, we used NGS Illumina mRNA-seq technology, which allowed us to avoid the technical issues inherent to microarray probe performance such as cross-hybridization, non-specific hybridization and limited detection range of individual probes (Agarwal *et al.*, 2010). By combining the higher resolution of mRNA-seq with an *in house* developed bioinformatics pipeline, we were able to find 300 intron retention, 79 alternative 3’ splice site usage, 48 Alt 5’ splice site usage, 8 ES and 350 multiple (complex) events (Fig. 3a) that are under circadian regulation. We also found 7 and 721 novel alternative exonic and intronic events, respectively. Interestingly, of the 646 genes that were found to contain cycling introns (Table S7), only 35 overlapped with the 499 reported by Hazen et al (7%, 35 out of 499).

In addition to documenting CCEs, we found that a clock-controlled splicing factor, AT2G02570, affected a subset of AS events when mutated. This gene had been previously identified as an SMN homolog (Kroiss *et al.*, 2008). SMN encodes the survival motor neuron protein, a spliceosome complex constituent which, when mutated, causes autosomal recessive spinal muscular atrophy (Talbot *et al.*, 1998; Kroiss *et al.*, 2008). However, our bioinformatics analysis revealed that AT2G02570 is more closely related to SMN’s paralogue SPF30 (Talbot *et al.*, 1998) (Fig. S8). Using the ASpli package, we identified 286 altered AS events in the AtSPF30 mutant *atspf30-1*. This is in accordance with previous reports that showed that changes in the levels or activities of core spliceosomal components or proteins that modulate the kinetics of spliceosome assembly can modulate alternative splicing (Saltzman *et al.*, 2011; Papasaikas *et al.*, 2015).

A comparison between the CCEs datasets and the *atspf30-1* dataset revealed that 32 events of the 286 events affected on an AtSPF30 mutant were among our PSI/PIR filtered subset of CCEs. Furthermore, a closer inspection of these splicing events showed that it preferentially affected IR events, by increasing overall retention level (Table S10). Indeed, we observed this effect upon validation of one of the shared events. qPCR analysis showed that, albeit with a similar pattern to what was observed for Col-0, IR of AT5G17670:E015 was higher at all assayed timepoints (Fig. 6f). This led us to believe that circadian control of AS is multifactorial because even though SPF30 is contributing by modulating the amplitude and overall level (i.e. higher IR), the underlying oscillation continues when SPF30 is absent.

In addition to AtSPF30, 30 other splicing related proteins were found to be under circadian regulation in all known circadian datasets (Fig. 4a). The contribution of each of these robust cycling factors on circadian control of AS remains to be determined and should be studied further. Prospectively, modulation of these circadian controlled splicing related factors could lead to time specific fine tuning of splicing events. Although many non-redundant splicing factors affect the clock when mutated (Sanchez *et al.*, 2010; Staiger & Green, 2011; Jones *et al.*, 2012; Wang *et al.*, 2012; Romanowski & Yanovsky, 2015; Schlaen *et al.*, 2015; Hernando *et al.*, 2017; Mateos *et al.*, 2018), we found no evidence that circadian modulation of splicing factors and related proteins led to AS changes in core clock components, but rather their effect was as an output (CCGs) mainly affecting other outputs (CCEs).

Splicing factors were also subject to circadian splicing events. In this sense, our stringent analysis revealed 57 splicing related genes that are also regulated at the splicing level (Table S7). A future careful and detailed study of the biological relevance of these splicing events will help us gain a better understanding of how the clock regulates alternative splicing, and vice versa.

It has been previously shown that core clock components are highly sensitive and can undergo AS upon environmental perturbations (Filichkin *et al.*, 2010; James *et al.*, 2012a; James *et al.*, 2012b; Syed *et al.*, 2012; Perez-Santangelo *et al.*, 2013; Staiger & Brown, 2013; Filichkin *et al.*, 2015; Mateos *et al.*, 2018). Furthermore, environmental disruptions appear to modulate AS of core clock components more than the clock itself. Taken together, this suggests that AS modulation of the central clock operates mainly via its input pathways (after an environmental perturbation), and circadian control of AS is mostly reserved to modulation of output pathways. However, we must point out that the number of CCEs reported in this work is very conservative and will likely increase in the future, as sequencing depth of circadian studies increases. The advent of novel long read technologies like those provided by Pacific Biosystems or Oxford Nanopore will likely allow for reliable transcript isoform level quantification at the global scale in the very near future.

## ACKNOWLEDGEMENTS

This work was supported by grants from Agencia Nacional de Promoción Científica y Tecnológica (ANPCyT) and the International Centre for Genetic Engineering and Biotechnology (ICGEB) to MY. AR was supported by postdoctoral fellowships from Comisión Nacional de Investigaciones Científicas (CONICET) and Fundación Bunge y Born (FBB). RGS was supported by Fundación Bunge y Born (FBB) and, SPS and EM were supported by CONICET.

We wish to thank Dr. Julieta Lisa Mateos and Philip Butlin for fruitful discussions and critical reading of the manuscript.

## AUTHOR CONTRIBUTIONS

AR, RGS, SPS and MY planned and designed the research; AR, RGS, SPS performed the experiments; AR, RGS and SPS analysed the experimental data; AR and EM performed the bioinformatics analysis, and AR and MY wrote the manuscript.

## SUPPORTING INFORMATION

**Fig. S1:** Correlation between RNA-seq samples and Multidimensional Scaling (MDS) Plot.

**Fig. S2:** Biological Process (BP) Gene Ontology (GO) analysis of clock-controlled genes (CCGs).

**Fig. S3:** Amplitude histogram of clock-controlled genes (CCGs) and core clock components.

**Fig. S4:** Biological Process (BP) Gene Ontology (GO) analysis of CCGs corresponding to the two largest phase groups.

**Fig. S5:** Comparison of genome coverage of RNA-seq, ATH1 microarray chip and AGRINOMICS1 technologies.

**Fig. S6:** CCGs are also clock regulated at the level of alternative use of exons and/or introns throughout the day.

**Fig. S7**: Validation of the COL2 IR splicing event, accompanying Fig. 3.

**Fig. S8:** Phylogenetic analysis of the splicing modulator AT2G02570.

**Table S1** Mapping statistics.

**Table S2** List of primers used in this work.

**Table S3** Rhythmic clock-controlled genes (CCGs) by JTK.

**Table S4** GO terms of CCGs (all).

**Table S5** Circadian regulated genes in different datasets.

**Table S6** GO terms of novel CCGs (all).

**Table S7** Rhythmic clock-controlled splicing events (CCEs) by JTK.

**Table S8** GO terms of genes associated to CCEs (all).

**Table S9** Splicing events misregulated in atspf30-1 mutant plants.

**Table S10** Misregulated events in atspf30-1 mutant plants that are also CCEs.

