## Supplementary material for "Global transcriptome analysis reveals circadian control of splicing events in *Arabidopsis thaliana*": Fig. S1 to S8 and Table S1-S2

The following Supporting Information is available for this article:

**Fig. S1** Correlation between RNA-seq samples and Multidimensional Scaling (MDS) Plot.

**Fig. S2** Biological Process (BP) Gene Ontology (GO) analysis of clock-controlled genes (CCGs).

**Fig. S3** Amplitude histogram of clock-controlled genes (CCGs) and core clock components.

**Fig. S4** Biological Process (BP) Gene Ontology (GO) analysis of CCGs corresponding to the two largest phase groups.

**Fig. S5** Comparison of genome coverage of RNA-seq, ATH1 microarray chip and AGRINOMICS1 technologies.

**Fig. S6** CCGs are also clock regulated at the level of alternative use of exons and/or introns throughout the day.

**Fig. S7** Validation of the COL2 IR splicing event, accompanying Fig. 3.

**Fig. S8** Phylogenetic analysis of the splicing modulator AT2G02570.

**Table S1** Mapping statistics.

**Table S2** List of primers used in this work.

**Table S3** Rhythmic clock-controlled genes (CCGs) by JTK.

**Table S4** GO terms of CCGs (all).

**Table S5** Circadian regulated genes in different datasets.

**Table S6** GO terms of novel CCGs (all).

**Table S7** Rhythmic clock-controlled splicing events (CCEs) by JTK.

**Table S8** GO terms of genes associated to CCEs (all).

**Table S9** Splicing events misregulated in atspf30-1 mutant plants.

**Table S10** Misregulated events in atspf30-1 mutant plants that are also CCEs.

**Fig. S1 Correlation between RNA-seq samples and Multidimensional Scaling (MDS) Plot.** Replicate correlation analysis of (A) LL 24 h, (B) LL 28 h, (C) LL 32 h, (D) LL 36 h, (E) LL 40 h, (F) LL 44 h, (G) LL 48 h, (H) LL 52 h, (I) LL 56 h, (J) LL 60h, (K) LL 64h, and (L) LL 68 h samples. (M) MDS plot of all the RNA-seq samples.

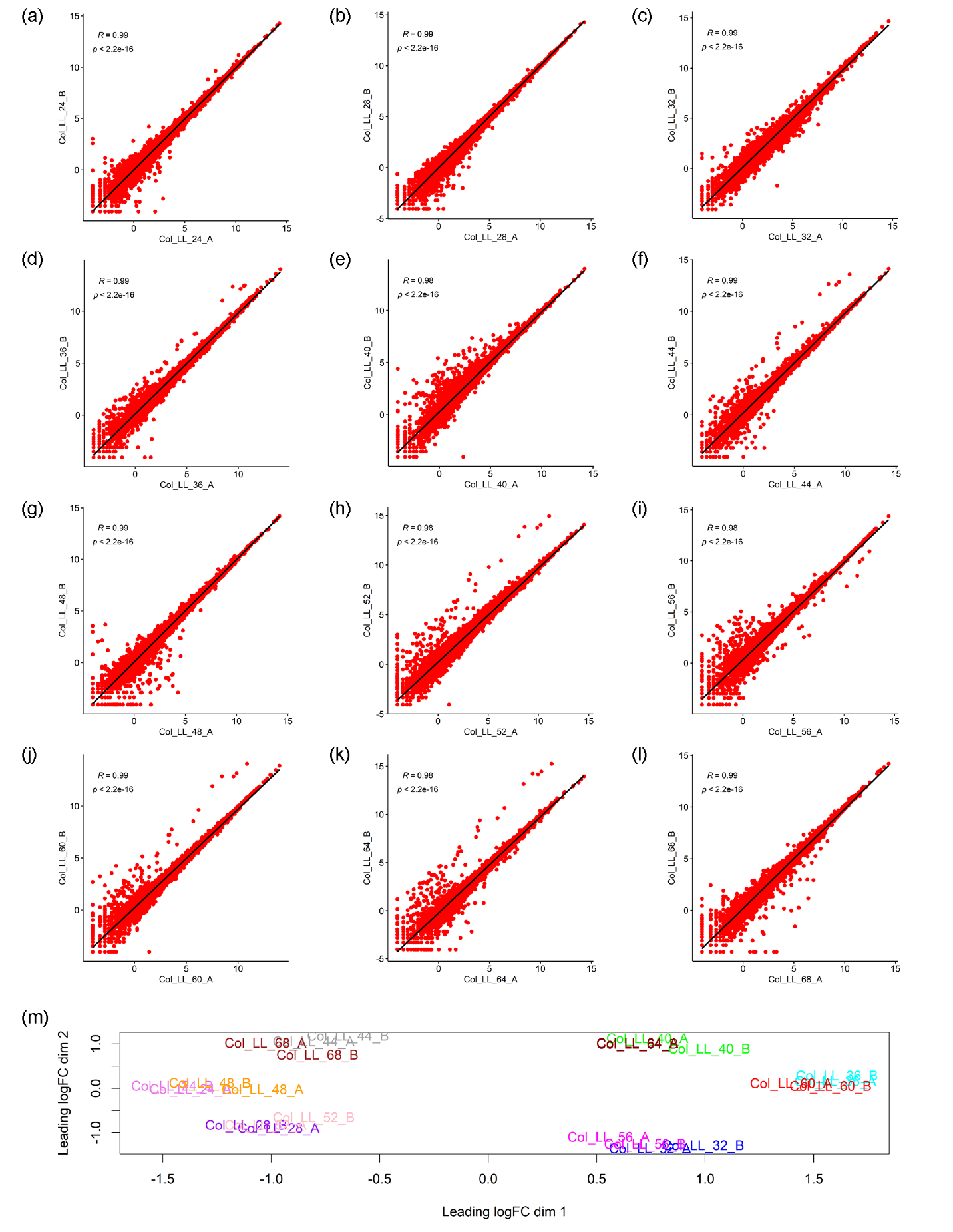

**Fig. S2 Biological Process (BP) Gene Ontology (GO) analysis of clock-controlled genes (CCGs).** Biological Process (BP) gene ontology analysis (GO) of all clock-controlled genes found in this study. Selected processes are shown. See Supplemental Table 4 for the full list.

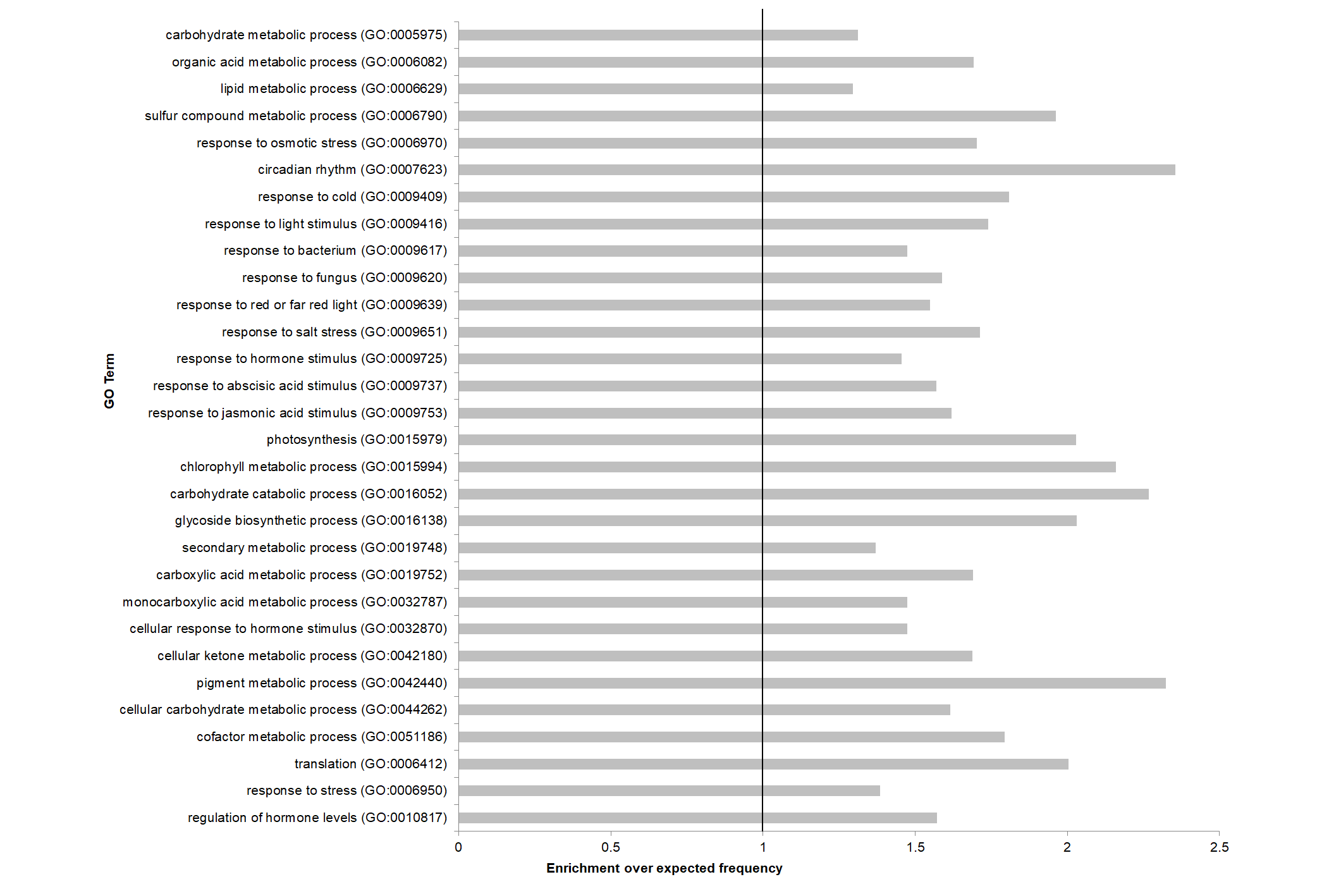

**Fig. S3 Amplitude histogram of clock-controlled genes (CCGs) and core clock components.** A histogram of the relative amplitude of al clock-controlled genes is shown. The inset shows the relative amplitude for core clock genes.

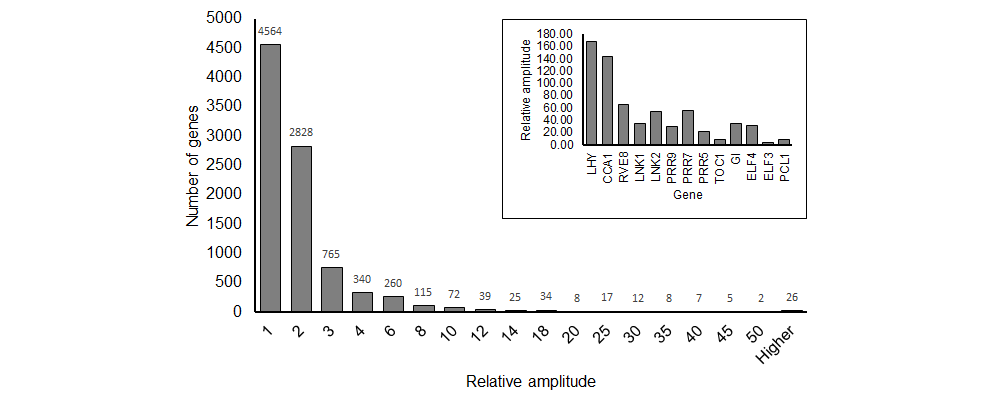

**Fig. S4 Biological Process (BP) Gene Ontology (GO) analysis of CCGs corresponding to the two largest phase groups.** Biological Process (BP) gene ontology analysis (GO) of the genes that exhibit a peak phase expression at LL10-12 (a) and 22-0 (b). Selected processes are shown. See Supplemental Table 6 for the full list.

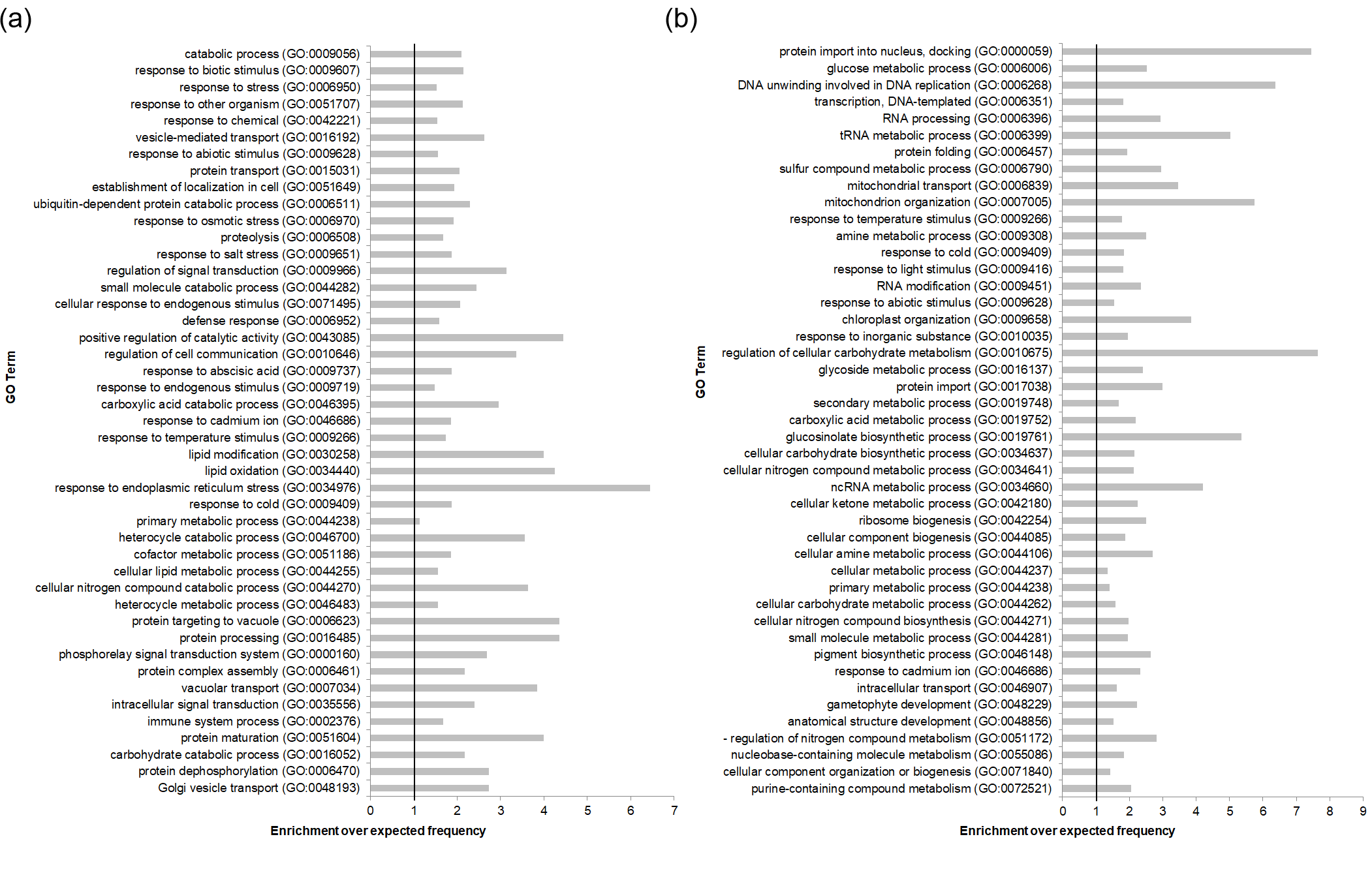

**Fig. S5 Comparison of genome coverage of RNA-seq, ATH1 microarray chip and AGRINOMICS1 technologies.** Total number of transcripts that can be detected with different global transcriptomics technologies. The table also shows the ratios of increase in transcript detection as compared to ATH1 and Agrinomics1.

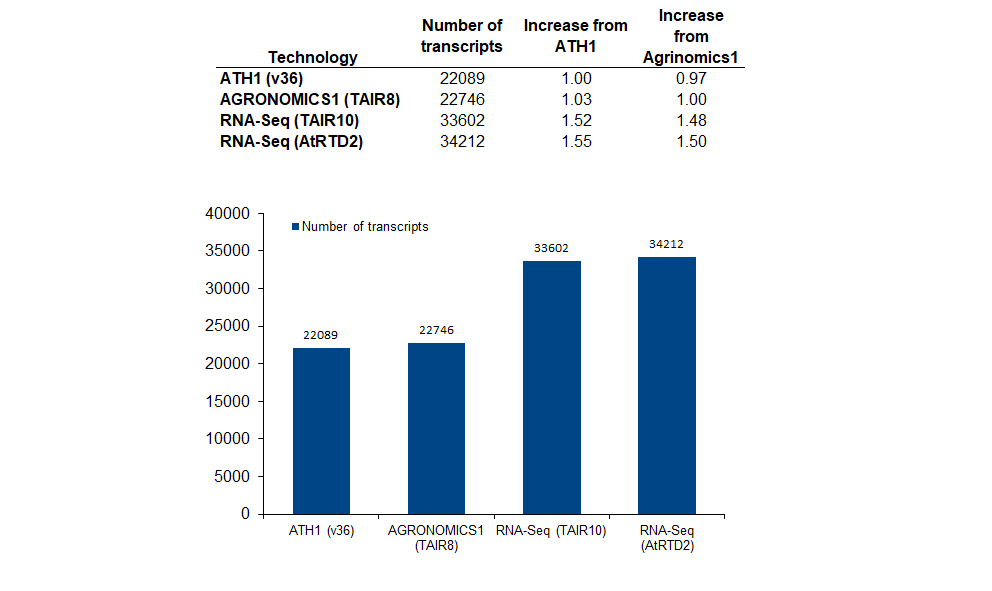

**Fig. S6 CCGs are also clock regulated at the level of alternative use of exons and/or introns throughout the day.** (a) Venn diagram comparing clock-controlled genes (CCGs) and genes which contain clock-controlled events (CCEs). (b) Number of genes detected in this work that underwent circadian control of alternative splicing at a novel, intron retention (IR), exon-skipping (ES), alt5’ss or alt3’ss level.

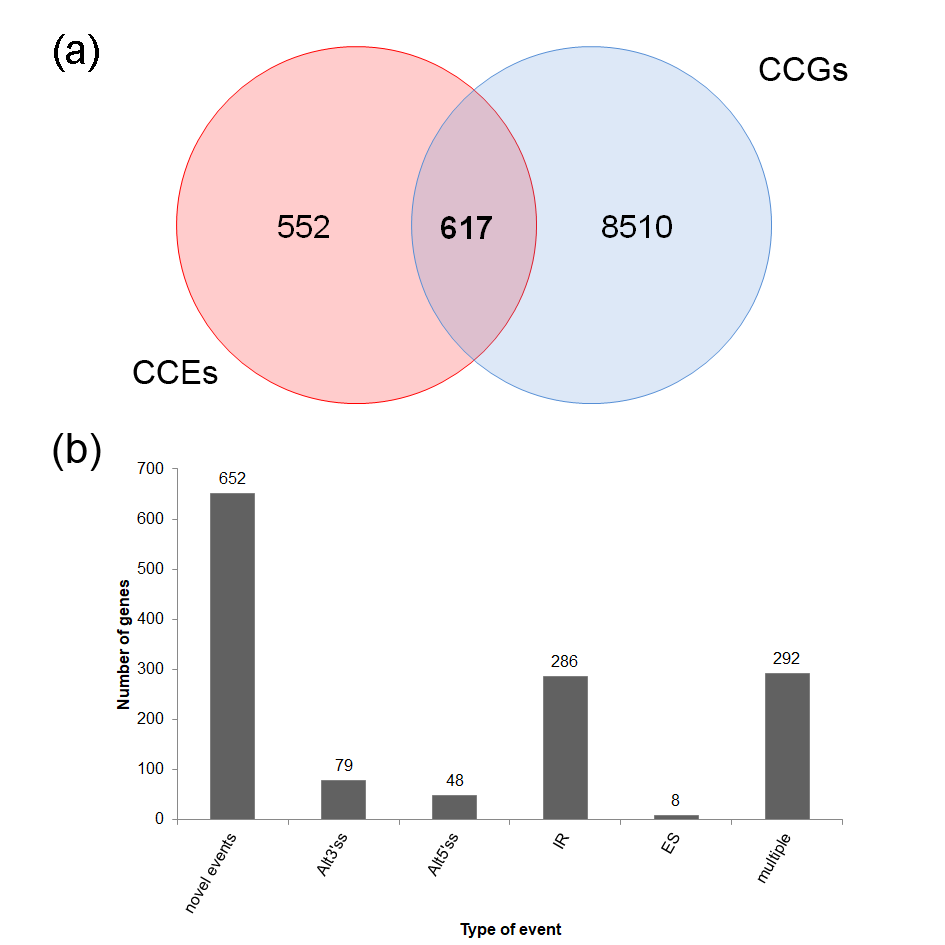

**Fig. S7 Validation of the COL2 IR splicing event, accompanying Fig. 3.** (a) Read density histogram of the intron retention (IR) event (highlighted in red) across the different samples (y axis). (b) RT-PCR gel agarose quantification of IR and full spliced isoforms the AT3G02380:E002 splicing event.

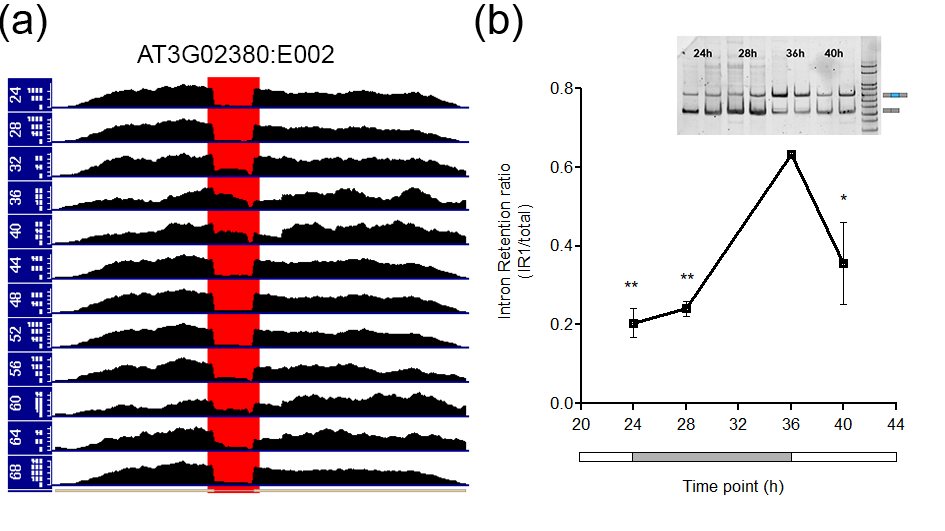

**Fig. S8 Phylogenetic analysis of the splicing modulator AT2G02570.** Phylogenetic analysis comparing metazoan SMN and SPF30 with homologs of the green lineage.

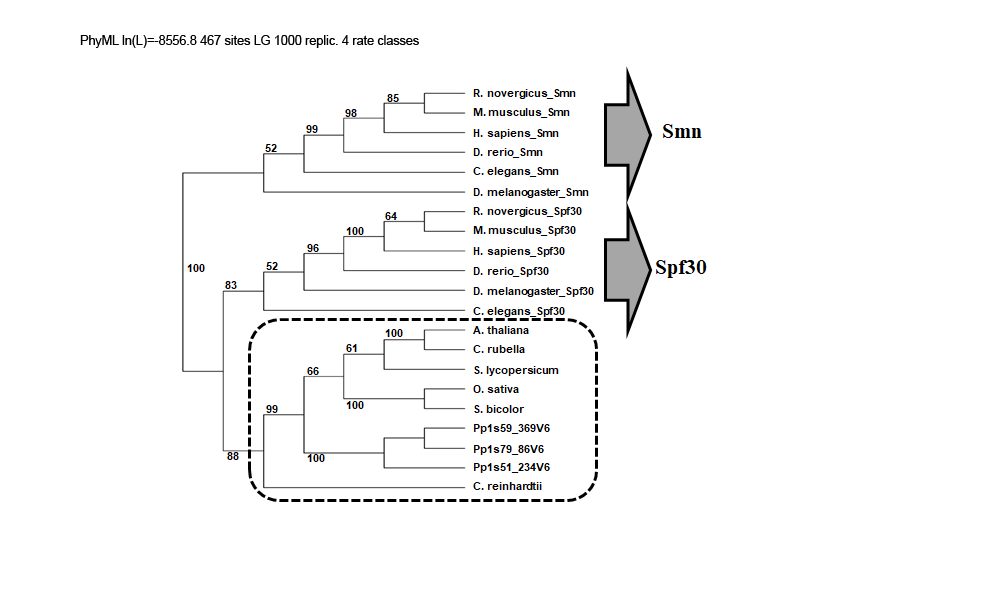

**Table S1** Mapping statistics.

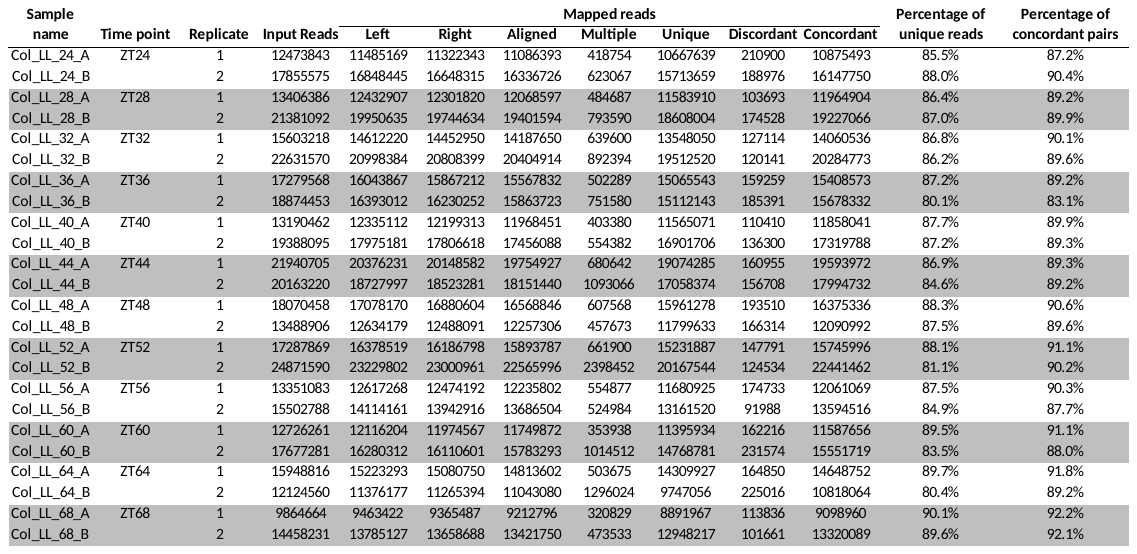

**Table S2** List of primers used in this work.

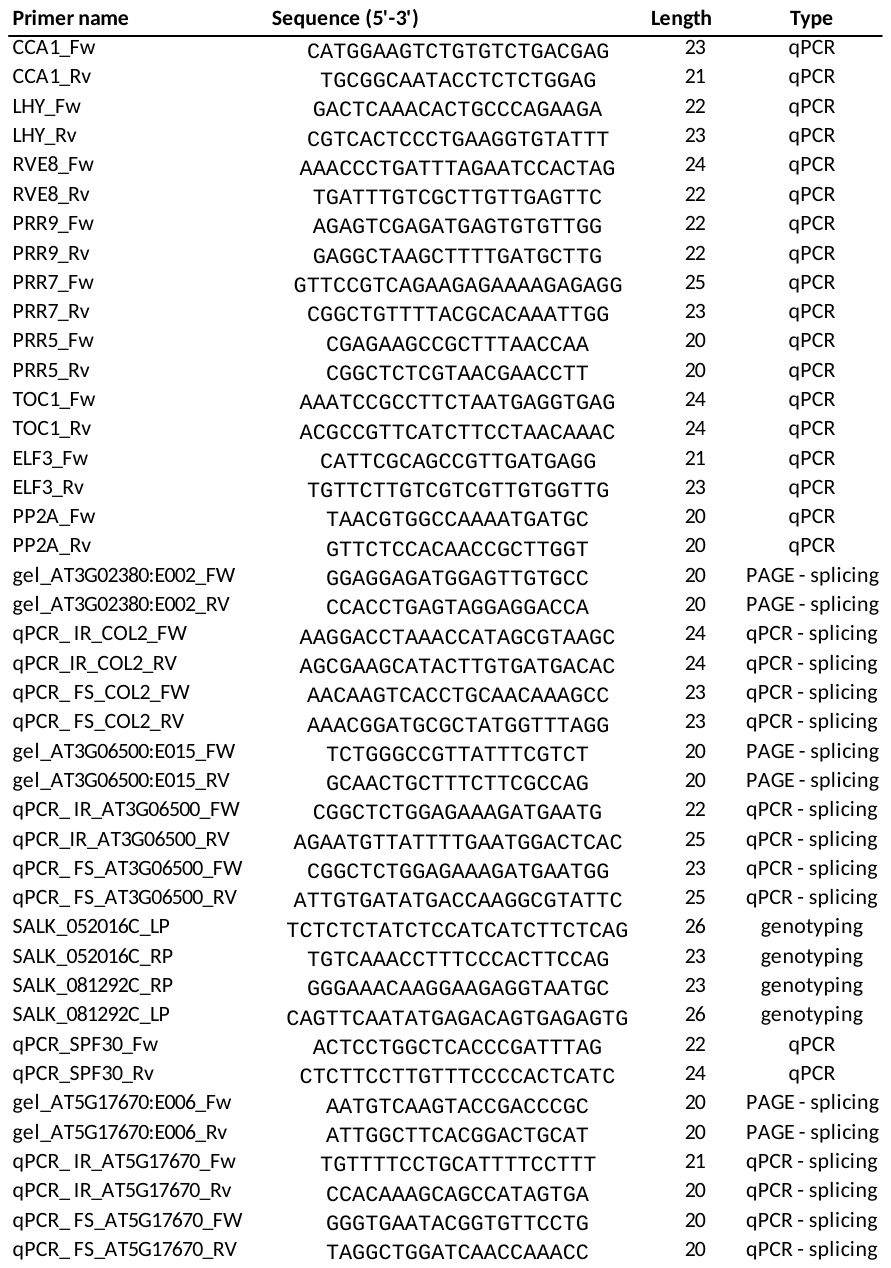
